# Gene model for the ortholog of *foxo* in *Drosophila sechellia*

**DOI:** 10.1101/2025.09.08.674750

**Authors:** Megan E. Lawson, Brandon A. Ruscoe, George G. Njuguna, Michelle L. Aguilar Medina, Sarah Raab, Eve Mellgren, Amy T. Hark, Natalie Minkovsky

## Abstract

Gene model for the ortholog of forkhead box, sub-group O (*foxo*) in the May 2011 (Broad dsec_caf1/DsecCAF1) Genome Assembly (GenBank Accession: GCA_000005215.1) of *Drosophila sechellia*. This ortholog was characterized as part of a developing dataset to study the evolution of the Insulin/insulin-like growth factor signaling pathway (IIS) across the genus *Drosophila* using the Genomics Education Partnership gene annotation protocol for Course-based Undergraduate Research Experiences.

## Introduction

*This article reports a predicted gene model generated by undergraduate work using a structured gene model annotation protocol defined by the Genomics Education Partnership (GEP; thegep.org) for Course-based Undergraduate Research Experience (CURE). The following information in quotes may be repeated in other articles submitted by participants using the same GEP CURE protocol for annotating Drosophila species orthologs of Drosophila melanogaster genes in the insulin signaling pathway*.

“In this GEP CURE protocol students use web-based tools to manually annotate genes in non-model *Drosophila* species based on orthology to genes in the well-annotated model organism fruitfly *Drosophila melanogaster*. The GEP uses web-based tools to allow undergraduates to participate in course-based research by generating manual annotations of genes in non-model species (Rele et al., 2023). Computational-based gene predictions in any organism are often improved by careful manual annotation and curation, allowing for more accurate analyses of gene and genome evolution (Mudge and Harrow 2016; Tello-Ruiz et al., 2019). These models of orthologous genes across species, such as the one presented here, then provide a reliable basis for further evolutionary genomic analyses when made available to the scientific community.” (Myers et al., 2024).

“The particular gene ortholog described here was characterized as part of a developing dataset to study the evolution of the Insulin/insulin-like growth factor signaling pathway (IIS) across the genus *Drosophila*. The Insulin/insulin-like growth factor signaling pathway (IIS) is a highly conserved signaling pathway in animals and is central to mediating organismal responses to nutrients (Hietakangas and Cohen 2009; Grewal 2009).” (Myers et al., 2024).

“A critical downstream effector of this pathway is the forkhead box O transcription factor (*foxo*, also designated FOXO1 or FKHR), which functions as a direct target of Akt kinase-mediated phosphorylation. Upon phosphorylation by Akt, FOXO undergoes cytoplasmic sequestration, resulting in downregulation of its transcriptional activity (Puig 2003). The *foxo* gene was originally identified in *Drosophila* through forward genetic screens that revealed gain-of-function mutations causing ectopic head-like structures in the foregut and hindgut regions (Jürgens 1988). Functional studies in fly larvae demonstrated that ectopic *foxo* expression recapitulates key aspects of the starvation response, including food avoidance behavior, developmental arrest, organ size reduction, and increased apoptosis (Kramer 2003; Jünger 2003).

Tissue-specific activation of *foxo* in the adult head fat body extends lifespan in both sexes and downregulates expression of insulin-like peptide 2 (*Ilp2*), establishing a regulatory feedback loop within the insulin signaling network (Hwangbo 2004). Conversely, loss-of-function *foxo* mutants exhibit reduced cell number without affecting individual cell size, indicating that *foxo* primarily regulates cell survival rather than cell growth (Jünger 2003).” (Kiser et al., 2025)

We propose a gene model for the *D. sechellia* ortholog of the *D. melanogaster* forkhead box, sub-group O (foxo) gene. The genomic region of the ortholog corresponds to the predicted forkhead box protein O LOC6606448 (RefSeq accession XP_032576567.1) in *D. sechellia* in the May 2011 (Broad dsec_caf1/DsecCAF1) Genome Assembly of *Drosophila sechellia* (GenBank Accession: GCA_000005215.1 - Drosophila 12 Genomes Consortium 2007). This gene model is based on RNA-Seq data from *D. sechellia* (McManus et al., 2014, Ma et al., 2018; PRJNA205470, PRJNA414017) and the foxo in *D. melanogaster* using FlyBase release FB2022_03 (Larkin et al., 2021; Gramates et al., 2022; Jenkins et al., 2022)

### Synteny

The target gene, *foxo*, occurs on chromosome 3R in *D. melanogaster* and is flanked upstream by *CG9922* and *CG3061*, and downstream by *Npc2b* and *CCHa1*. The tblastn search of *D. melanogaster* foxo-RB_prot (query) against the *D. sechellia* (GenBank Accession: GCA_ GCA_000005215.1) Genome Assembly (database) placed the putative ortholog of f*oxo* within scaffold super_0 (CH480815.1) at locus LOC6606448 (XP_032576567.1)— with an E-value of 0.0 and a percent identity of 95.53%. Furthermore, the putative ortholog is flanked upstream by LOC6606450 (XP_002031253.1) and LOC6606451 (XP_002031254.1), which correspond to *CG9922* and *CG3061* in *D. melanogaster* (E-value: 1e-57 and 0.0; identity: 99.19% and 95.68%, respectively, as determined by blastp; Figure 1A, Altschul et al., 1990). The putative ortholog of *foxo* is flanked downstream by LOC6606447 (XP_032577736.1) and LOC6606446 (XP_002031250.1), which correspond to *Npc2b* and *CCHa1* in *D. melanogaster* (E-value: 1e-115 and 5e-47; identity: 98.74% and 87.36%, respectively, as determined by blastp). The putative ortholog assignment for *foxo* in *D. sechellia* is supported by the following evidence: The genes surrounding the *foxo* ortholog are orthologous to the genes at the same locus in *D. melanogaster* and local synteny is completely conserved, supported by high quality matches generated from *blastp*; we conclude that LOC6606448 is the correct ortholog of *foxo* in *D. sechellia* (Figure 1A).

**Figure 1:**
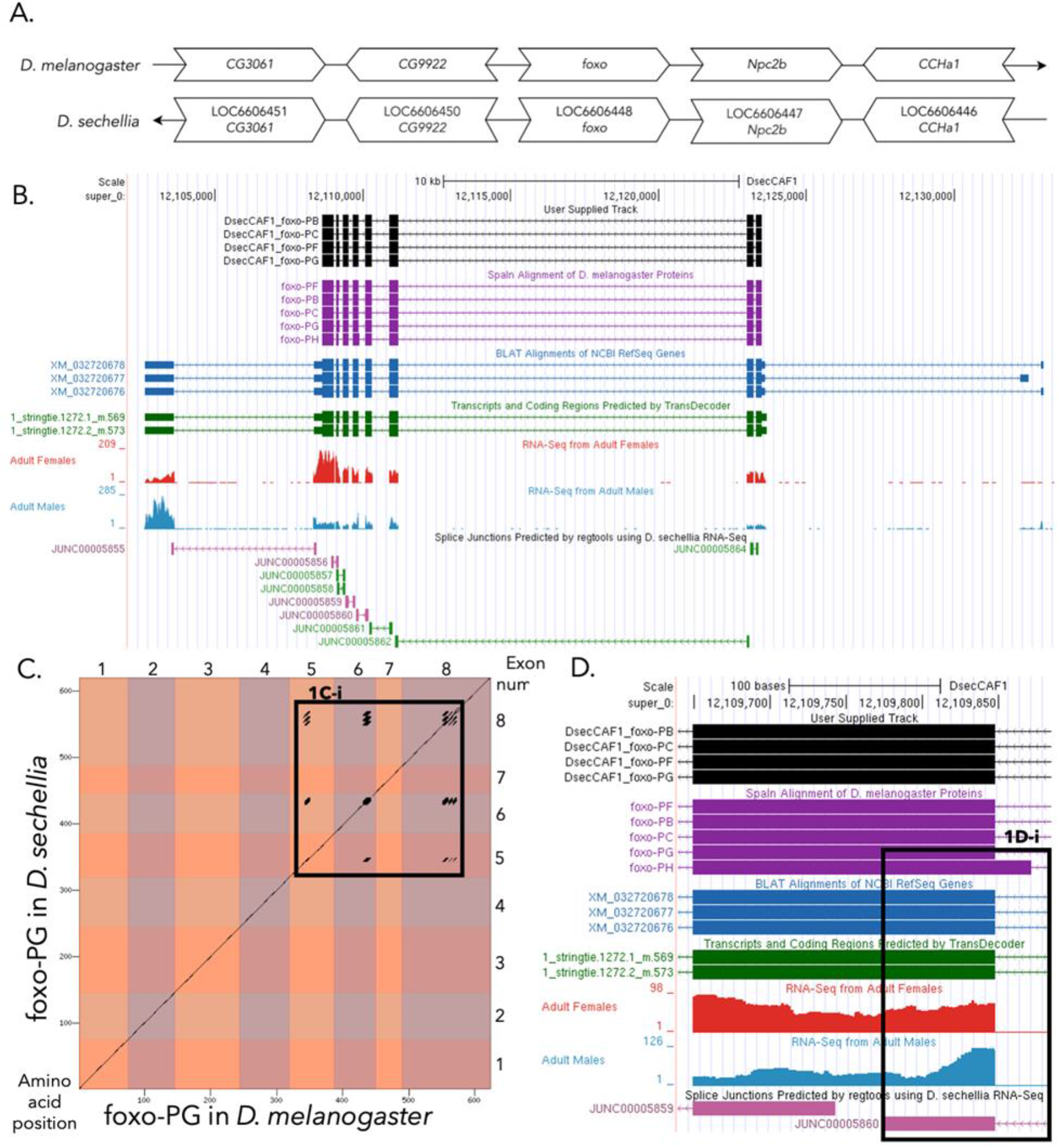
*foxo* gene model comparison between *Drosophila sechellia* and *Drosophila melanogaster* orthologs. **(A) Synteny comparison of the genomic neighborhoods for *foxo* in *Drosophila melanogaster* and *D. sechellia***. Thin arrows indicate the DNA strand within which the target gene– *foxo* –is located in *D. melanogaster* (top) and *D. sechellia* (bottom). The thin arrow pointing to the left indicates that *foxo* is on the negative (-) strand in *D. sechellia*, and the thin arrow pointing to the right indicates that foxo is on the positive (+) strand in D. melanogaster. The wide gene arrows pointing in the same direction as foxo are on the same strand relative to the thin arrows, while wide gene arrows pointing in the opposite direction of foxo are on the opposite strand relative to the thin arrows. White gene arrows in *D. sechellia* indicate orthology to the corresponding gene in *D. melanogaster*. Gene symbols given in the *D. sechellia* gene arrows indicate the orthologous gene in *D. melanogaster*, while the locus identifiers are specific to *D. sechellia* **(B) Gene Model in GEP UCSC Track Data Hub (Raney et al**., **2014)**. The coding-regions of *foxo* in *D. sechellia* are displayed in the User Supplied Track (black); coding CDSs are depicted by thick rectangles and introns by thin lines with arrows indicating the direction of transcription. Subsequent evidence tracks include BLAT Alignments of NCBI RefSeq Genes (dark blue, alignment of RefSeq genes for *D. sechellia*), Spaln of *D. melanogaster* Proteins (purple, alignment of Ref-Seq proteins from *D. melanogaster*), Transcripts and Coding Regions Predicted by TransDecoder (dark green), RNA-Seq from Adult Females and Adult Males (red and light blue respectively; alignment of Illumina RNA-Seq reads from *D. sechellia*), Splice Junctions Predicted by regtools using *D. sechellia* RNA-Seq (McManus et al., 2014, Ma et al., 2018; PRJNA205470, PRJNA414017). The splice junctions pertaining to the *foxo* ortholog, have a minimum read-depth of 55 with 50-99 and 100-499 supporting reads in green and pink respectively. **(C) Dot Plot of foxo-PG in *D. melanogaster* (x-axis) vs. the orthologous peptide in *D. sechellia* (y-axis)**. Amino acid number is indicated along the left and bottom; CDS number is indicated along the top and right, and CDSs are also highlighted with alternating colors. Line breaks in the dot plot indicate mismatching amino acids at the specified location between species. The box in figure C (1C-i) shows a region of LINE repeats in the protein sequence. (D) Model in UCSC Track Hub demonstrating the lack of data to support the existence of *foxo*-PH isoform in *D. sechellia* (Raney et al., 2014).

### Protein Model

*foxo* in *D. sechellia* has four protein coding isoforms, foxo-PB, foxo-PC, foxo-PF, and foxo-PG (Figure 1B). Isoforms foxo-PB and foxo-PC are identical and contain eight CDS. Isoforms foxo-PF and foxo-PG are identical and contain eight CDS. All these isoforms are also present in the *foxo* ortholog in *D. melanogaster*, each containing the same number of exons across both species. However, *foxo* in *D. melanogaster* has a fifth isoform, foxo-PH, which is not present in *D. sechellia*. Isoform foxo-PH also contains eight CDS. The absence of this isoform in *D. sechellia* was determined due to the lack of splice junctions, RNA-Seq data, and a canonical acceptor site that would maintain the correct reading frame (Figure D). The sequence of foxo-PG in *D. sechellia* has 97.3% identity with isoform foxo-PG in *D. melanogaster* as determined by *blastp* (Figure 1C).

### Special characteristics of the protein model

Isoform foxo-PH in *D. melanogaster* is differentiated from isoforms foxo-PB and foxo-PC by a unique splice site at the beginning of the fifth exon. However, there are no alternative predicted splice junctions, no canonical acceptor sites that would maintain the correct reading frame, and no RNA-Seq data in this region to support the existence of this unique splice site (Figure D). This evidence indicates that foxo-PH is likely not present in *D. sechellia*.

## Methods

Detailed methods including algorithms, database versions, and citations for the complete annotation process can be found in Rele et al. (2023). Briefly, students use the GEP instance of the UCSC Genome Browser v.435 (https://gander.wustl.edu; Kent WJ et al., 2002; Navarro Gonzalez et al., 2021) to examine the genomic neighborhood of their reference IIS gene in the *D. melanogaster* genome assembly (Aug. 2014; BDGP Release 6 + ISO1 MT/dm6). Students then retrieve the protein sequence for the *D. melanogaster* reference gene for a given isoform and run it using *tblastn* against their target *Drosophila* species genome assembly on the NCBI BLAST server (https://blast.ncbi.nlm.nih.gov/Blast.cgi; Altschul et al., 1990) to identify potential orthologs. To validate the potential ortholog, students compare the local genomic neighborhood of their potential ortholog with the genomic neighborhood of their reference gene in *D. melanogaster*. This local synteny analysis includes at minimum the two upstream and downstream genes relative to their putative ortholog. They also explore other sets of genomic evidence using multiple alignment tracks in the Genome Browser, including BLAT alignments of RefSeq Genes, Spaln alignment of *D. melanogaster* proteins, multiple gene prediction tracks (e.g., GeMoMa, Geneid, Augustus), and modENCODE RNA-Seq from the target species. Detailed explanation of how these lines of genomic evidenced are leveraged by students in gene model development are described in Rele et al. (2023). Genomic structure information (e.g., CDSs, intron-exon number and boundaries, number of isoforms) for the *D. melanogaster* reference gene is retrieved through the Gene Record Finder (https://gander.wustl.edu/~wilson/dmelgenerecord/index.html; Rele et al., 2023). Approximate splice sites within the target gene are determined using *tblastn* using the CDSs from the *D. melanogaste*r reference gene. Coordinates of CDSs are then refined by examining aligned modENCODE RNA-Seq data, and by applying paradigms of molecular biology such as identifying canonical splice site sequences and ensuring the maintenance of an open reading frame across hypothesized splice sites. Students then confirm the biological validity of their target gene model using the Gene Model Checker (https://gander.wustl.edu/~wilson/dmelgenerecord/index.html; Rele et al., 2023), which compares the structure and translated sequence from their hypothesized target gene model against the *D. melanogaster* reference gene model. At least two independent models for a gene are generated by students under mentorship of their faculty course instructors. Those models are then reconciled by a third independent researcher mentored by the project leaders to produce the final model. Note: comparison of 5’ and 3’ UTR sequence information is not included in this GEP CURE protocol. (Lawson et al., 2025)

## Supporting information

Gene model data files

## Supplemental Files

1. Zip file containing a FASTA, PEP, GFF files for the gene model

2. Figure 1 in high resolution

## Acknowledgements

We would like to thank Wilson Leung for developing and maintaining the technological infrastructure that was used to create this gene model and Laura K. Reed for overseeing the project. Thank you to FlyBase for providing the definitive database for *Drosophila melanogaster* gene models. FlyBase is supported by grants: NHGRI U41HG000739 and U24HG010859, UK Medical Research Council MR/W024233/1, NSF 2035515 and 2039324, BBSRC BB/T014008/1, and Wellcome Trust PLM13398.

